# Trimmed Constrained Mixed Effects Models: Formulations and Algorithms

**DOI:** 10.1101/2020.01.28.923599

**Authors:** Peng Zheng, Ryan Barber, Reed Sorensen, Christopher Murray, Aleksandr Aravkin

## Abstract

Mixed effects (ME) models inform a vast array of problems in the physical and social sciences, and are pervasive in meta-analysis. We consider ME models where the random effects component is linear. We then develop an efficient approach for a broad problem class that allows nonlinear measurements, priors, and constraints, and finds robust estimates in all of these cases using trimming in the associated marginal likelihood.

The software accompanying this paper is disseminated as an open-source Python package called `LimeTr`. `LimeTr` is able to recover results more accurately in the presence of outliers compared to available packages for both standard longitudinal analysis and meta-analysis, and is also more computationally efficient than competing robust alternatives. Supplementary materials that reproduce the simulations, as well as run `LimeTr` and third party code are available online. We also present analyses of global health data, where we use advanced functionality of `LimeTr`, including constraints to impose monotonicity and concavity for dose-response relationships. Nonlinear observation models allow new analyses in place of classic approximations, such as log-linear models. Robust extensions in all analyses ensure that spurious data points do not drive our understanding of either mean relationships or between-study heterogeneity.

## 1 Introduction

Linear mixed effects (LME) models play a central role in a wide range of analyses [Bates et al., 2015]. Examples include longitudinal analysis [Laird et al., 1982], meta-analysis [DerSimonian and Laird, 1986], and numerous domain-specific applications [Zuur et al., 2009].

Robust LME models are typically obtained by using heavy tailed error models for random effects. The Student’s t distribution [Pinheiro et al., 2001], as well as weighting functions [Koller, 2016] have been used. The resulting formulations are fit either by EM methods, estimating equations, or by MCMC [Rosa et al., 2003]. In this paper, we take a different track, and extend the least trimmed squares (LTS) method to the ME setting. While LTS has found wide use in a range of applications [Aravkin and Davis, 2019, Yang and Lozano, 2015, Yang et al., 2018], trimming the ME likelihood extends prior work.

### Contributions

In this paper, we consider a subclass of nonlinear mixed effects models. We allow nonlinear measurements, priors, and constraints, but require that the random effects enter the model in a linear way. We call this class *partially nonlinear* ME models, and it covers a broad class of problems while allowing tractable algorithms. We develop new conditions that guarantee the existence of estimators for partially nonlinear models, a trimming approach that robustifies any linear or partially nonlinear model against outliers, and algorithms for solving the nonconvex optimization problems required to find estimates with standard guarantees (convergence to stationary points). We also show splines (and associated shape constraints) can be used to capture key nonlinear relationships, and illustrate the full modeling capability on real-data examples based on dose-response relationships.

The main code to perform the inference is published as open source Python package called `LimeTr` (Linear Mixed Effects with Trimming, pronounced *lime tree*). All synthetic experiments using `LimeTr` have been submitted for review as supplementary material with this paper. The `LimeTr` package allows functionality that is not available through other available open source tools. The functionality of `LimeTr` is summarized in Table 1.

**Table 1:**
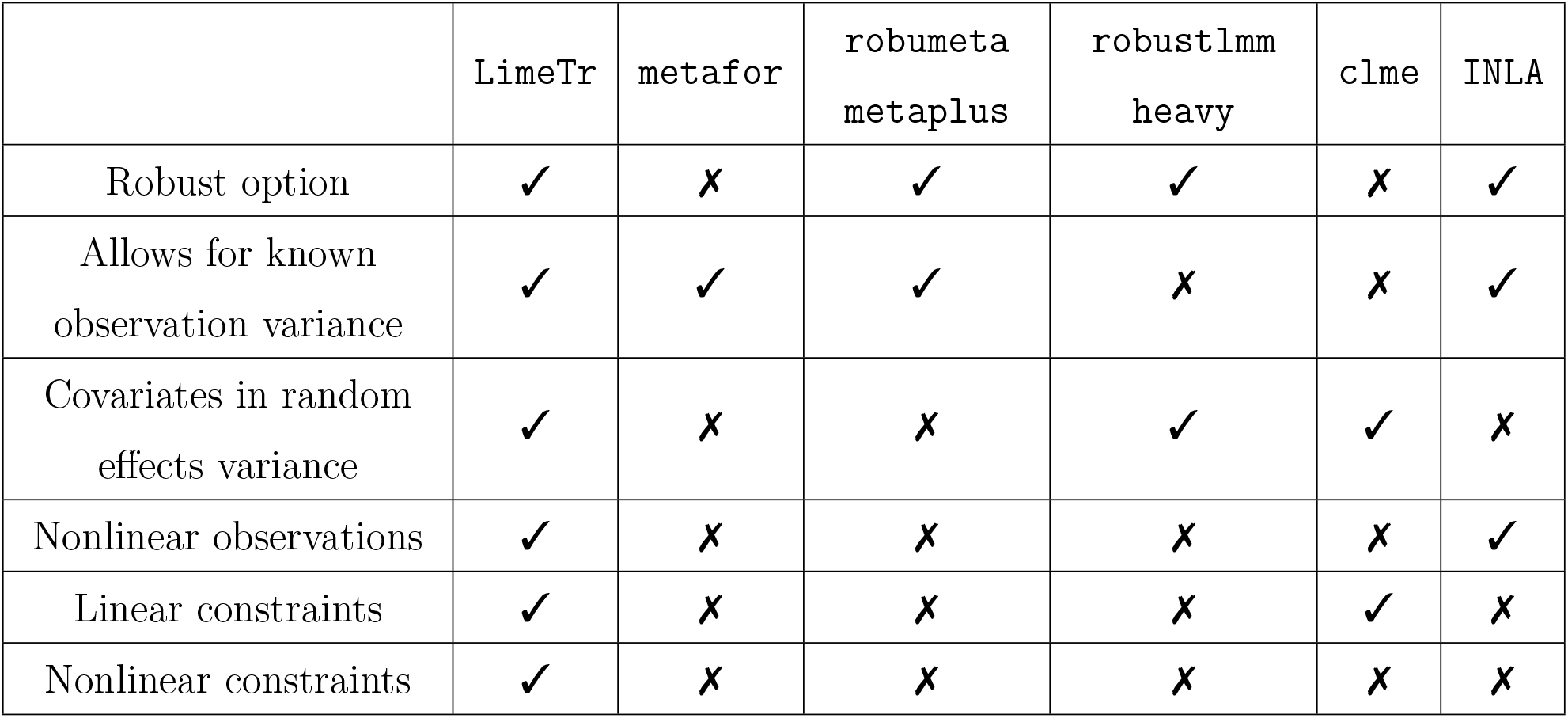
Comparison with currently available robust mixed effects packages.

The paper proceeds as follows. In Section 2.2, we describe the problem class of ME models and derive the marginal maximum likelihood (ML) estimator. In Section 2.3, we describe how constraints and priors are imposed on parameters. In Section 2.4, we review trimming approaches and develop a new trimming extension for the ML approach. In Section 2.5, we present a customized algorithm based on variable projection, along with a convergence analysis. In Section 2.6, we discuss spline models for dose-response relationships and give examples of shape-constrained trimmed spline models. Section 3 shows the efficacy of the methods for synthetic and empirical data. In Section 3.1, we validate the ability of the method to detect outliers when working with heterogeneous longitudinal data, and compare with other packages. In Section 3.2 we apply the method to analyze empirical data sets for both linear and nonlinear relationships using trimmed constrained MEs. This section highlights new capability of `LimeTr` that is not available in other packages.

## 2 Methods

### 2.1 Notation and Modeling Concepts

In this section, we define notation and concepts used throughout the paper. Additional definitions and notation are introduced in the analysis section. We use lower case letters to denote scalars, e.g. *β*, and scalar-valued functions, e.g. *f* (*β*), and bold letters represent vectors, e.g. ***β***, and vector-valued functions, e.g. ***f***_*i*_(***β***). We use capital bold letters to represent matrices, e.g. ***X***. All variables and vectors are real, i.e. in ℝ^*k*^, with *k* indicating dimension. For a smooth function *f*: ℝ^*k*^ → ℝ, we denote the vector of first derivatives, or **gradient**, by ∇*f*, and the matrix of second derivatives, or **Hessian**, by ∇^2^*f*. We use diag(***x***) to denote a diagonal matrix whose diagonal is an input vector ***x***. We use ***M*** ^*−*1^ to denote the inverse of a matrix, and denote weighted norms (Mahalanobis distances) by

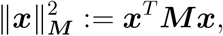

and |**M**| to denote the determinant of **M**. We use the ⊙ notation to denote the Hadamard product or operation, so in particular

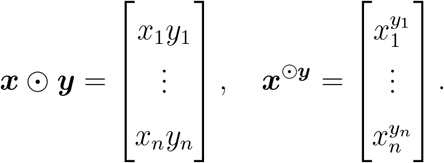

A **likelihood** maps parameters to an associated density function for observed data. In the mixed effects context, these parameters can be separated into **fixed effects** (e.g. population mean) and **random effects** (e.g. study-specific random intercept). A **marginal** likelihood function refers to the likelihood obtained by integrating out random effects from the joint likelihood.

We incorporate additional information about parameters using statistical priors, and restrict parameter domains using **constraints**. When constraints are present we use the term **constrained likelihood**. We use **trimming** (Section 2.4) to robustify a (marginal) likelihood, and use the term **trimmed likelihood** to describe such likelihoods.

The goal of an inference problem is to maximize the (marginal) likelihood or modified likelihood, or equivalently to minimize the negative logarithm of such a function, called an **objective function** in optimization. The optimization problem specification includes constraints as well as the objective. We use min to refer to the minimum value of an optimization problem, and arg min to refer to the minimizer, which corresponds to the **estimator** in this setting.

We define projection of a point ***x*** onto a closed set *C* by

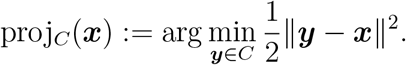

### 2.2 Problem Class

We consider the following mixed effects model:

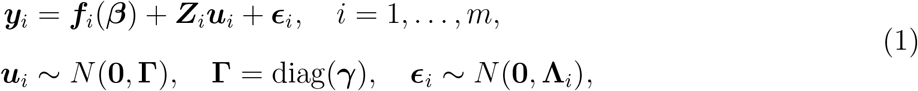

where *m* is the number of groups, 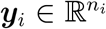 is the vector of observations from the *i*th group, and *n* =∑_*i*_ *n*_*i*_ is the total number of observations. Measurement errors are denoted by 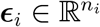 with covariance 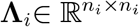, and we denote by **Λ** ∈ ℝ^*n×n*^ the full block diagonal measurement error covariance matrix. Regression coefficients are denoted by 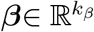. Random effects are denoted by 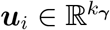, where 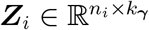 are linear maps. The functions 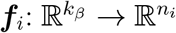 may be nonlinear, but we restrict the random effects to enter in a linear way through the linear maps 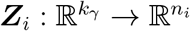.

A range of assumptions may be placed on **Λ**. In longitudinal analysis, **Λ** is often a diagonal or block-diagonal matrix, parametrized by a small set of parameters, with the simplest example **Λ** = *σ*^2^***I*** where *σ*^2^ is unknown. In meta-analysis, **Λ** is a known diagonal matrix whose entries are variances for each input datum. We do not restrict the term ‘meta-analysis’ to a single observation per study, since many analyses include multiple observations, such as summary results for quartiles based on exposure. The distinguishing feature of meta-analysis is the specification of a known **Λ** matrix.

For convenience, we denote by ***θ*** the tuple of fixed parameters:

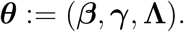

The joint likelihood corresponding to model (1) for ***θ*** and random effects ***u*** is given by

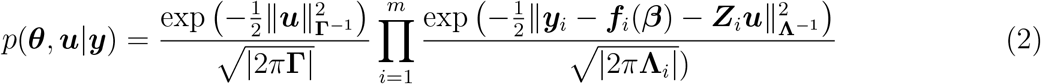

Maximizing (2) with respect to both fixed and random parameters is problematic, as the number of random parameters ***u***_*i*_ grows with the number of groups. In the extreme case of one observation per group, there are more unknowns than datapoints. Standard practice is to marginalize random effects, integrating (2) with respect to all ***u***_*i*_. The numerical estimation is then accomplished by taking the negative logarithm (a simplifying transformation) and minimizing the result, which is an equivalent problem to maximizing the marginal likelihood:

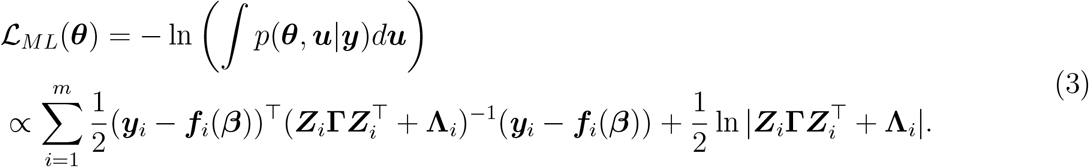

Problem (3) is equivalent to a maximum likelihood formulation arising from a Gaussian model with correlated errors:

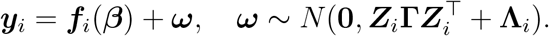

The structure of this objective depends on the structural assumptions on **Λ**. In the scope of this paper we always assume **Λ** is a diagonal matrix, namely all the measurement error are independent with each other. We restrict our numerical experiments to two particular classes: (1) **Λ** = *σ*^2^***I*** with *σ*^2^ unknown, used in standard longitudinal analysis, and (2) the measurements variance is provided and used in meta-analysis. `LimeTr` allows other structure options as well, for example group-specific unknown 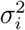 that extends case (1), but we consider a simple set of synthetic results to help focus on robust capabilities.

### 2.3 Constraints and Priors

The ML estimate (3) can be extended to incorporate linear and nonlinear inequality constraints

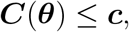

where ***θ*** are any parameters of interest. Constraints play a key role in Section 2.6, when we use polynomial splines to model nonlinear relationships. The trimming approach developed in the next section is applicable to both constrained and unconstrained ML estimates.

In many applications it is essential to allow priors on parameters of interest ***θ***. We assume that priors follow a distribution defined by the density function

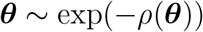

where *ρ* is smooth (but may be nonlinear and nonconvex). The likelihood problem is then augmented by adding the term *ρ*(***θ***) to the ML objective (3). The most common use case is 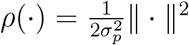, for some user-defined 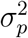.

In the next section we describe trimmed estimators, and extend them to the ME setting.

### 2.4 Trimming in Mixed Effect Models

Least trimmed squares (LTS) is a robust estimator proposed by Rousseeuw [1985], Rousseeuw and Croux [1993] for the standard regression problem. Starting from a standard least squares estimator,

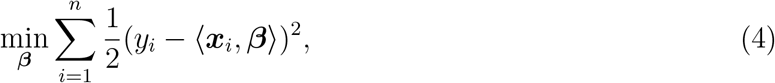

the LTS approach modifies (4) to minimize the sum of *smallest h* residuals rather than all residuals. These estimators were initially introduced to develop linear regression estimators that have a high breakdown point (in this case 50%) and good statistical efficiency (in this case *n*^*−*1*/*2^).^1^ LTS estimators are robust against outliers, and arbitrarily large deviations that are trimmed do not affect the final estimate.

The explicit LTS extension to (4) is formed by introducing auxiliary variables ***w***:

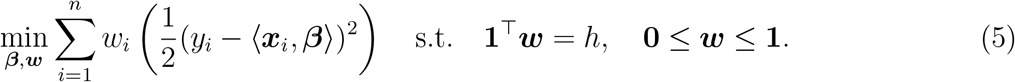

The set

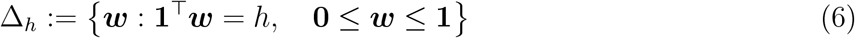

is known as the *capped simplex*, since it is the intersection of the *h*-simplex with the unit box (see e.g. Aravkin and Davis [2019] for details). For a fixed ***β***, the optimal solution of (5) with respect to ***w*** assigns weight 1 to each of the smallest *h* residuals, and 0 to the rest. Problem (5) is solved *jointly* in (***β, w***), simultaneously finding the regression estimate and classifying the observations into inliers and outliers. This joint strategy makes LTS different from post hoc analysis, where a model is first fit with all data, and then outliers are detected using that estimate.

Several approaches for finding LTS and other trimmed M-estimators have been developed, including FAST-LTS [Rousseeuw and Van Driessen, 2006], and exact algorithms with exponential complexity [Mount et al., 2014]. The LTS approach (5) does not depend on the form of the least squares function, and this insight has been used to extend LTS to a broad range of estimation problems, including generalized linear models [Neykov and Müller, 2003], high dimensional sparse regression [Alfons et al., 2013], and graphical lasso [Yang and Lozano, 2015, Yang et al., 2018].

The most general problem class to date, presented by Aravkin and Davis [2019], is formulated as

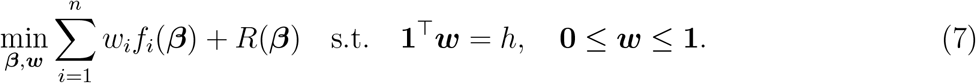

where *f*_*i*_ are continuously differentiable (possibly nonconvex) functions and *R* describes any regularizers and constraints (which may also be nonconvex).

Problem (3) is not of the form (7), except for the special case where we want to detect *entire outlying groups*. This is limiting, since we want to differentiate measurements within groups. We solve the problem by using a new trimming formulation.

To explain the approach we focus on trimming a single group term from the likelihood (3):

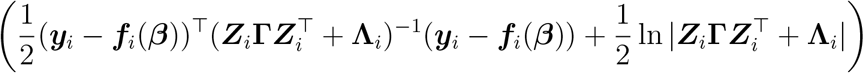

Here, 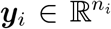, where *n*_*i*_ is the number of observations in the *i*th group. To trim observations within the group, we introduce auxiliary variables 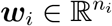, and define

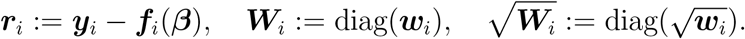

We now form the objective

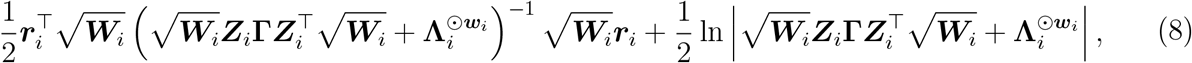

where ⨀ denotes the elementwise power operation:

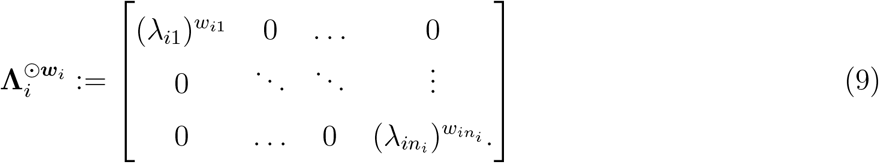

When *w*_*ij*_ = 1, we recover the contribution of the *ij*th observation to the original likelihood. As *w*_*ij*_ ↓ 0, The *ij*th contribution to the residual is correctly eliminated by 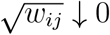. The *j*th row and column of 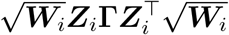 both go to 0, while the *j*th entry of 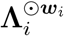 goes to 1, which removes all impact of the *j*th point. Specifically, the matrix 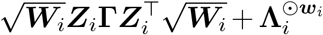 with *w*_*ij*_ = 0 and all remaining *w*_*i·*_ = 1 is the same as the matrix 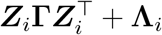 obtained after deleting the *ij*-th point.

Combining trimmed ML with priors and constraints, we obtain the following modified log-likelihood:

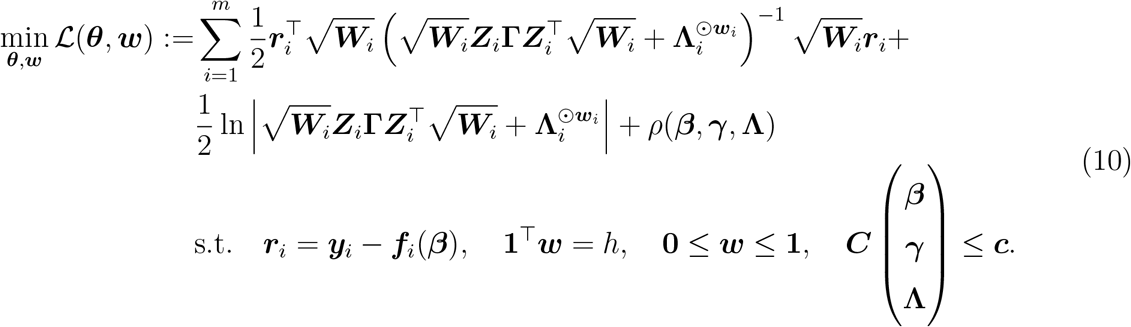

Problem (10) has not been previously considered in the literature. We present a specialized algorithm and analysis in the next section.

### 2.5 Fitting Trimmed Constrained MEs: Algorithm and Analysis

Problem (10) is nonsmooth and nonconvex. The key to algorithm design and analysis is to decouple this structure, and reduce the estimator to solving a smooth nonconvex value function over a convex set. This allows an efficient approach that combines classic nonlinear programming with first-order approaches for optimizing nonsmooth nonconvex problems. We partially minimize with respect to (***β, γ*, Λ**) using an interior point method, and then optimize the resulting value function with respect to ***w*** using a first-order method. The approach leverages ideas from variable projection [Golub and Pereyra, 1973, 2003, Aravkin and Van Leeuwen, 2012, Aravkin et al., 2018].

We define ***θ*** = (***β, γ*, Λ**), the implicit solution ***θ***(***w***) and value function *v*(***w***) as follows:

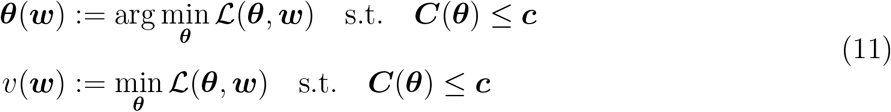

where ℒ(***θ, w***) is given in (10). The term ***θ***(***w***) refers to the entire set of minimizers for a given *w*, which may not necessarily be a singleton.

We first develop conditions and theory to guarantee the existence of minimizers ***θ***(***w***). While there are results in the literature for particular classes of linear mixed effects models, conditions for the existence of minimizers for the partial nonlinear case (10) have not been derived. Both Harville [2018] and Davidian and Giltinan [1995] analyze the Aitken model, where **Λ** = *σ*^2^***H*** for a nonsingular ***H***, by essentially deriving the closed form estimates for ***θ*** in this case, where the conditions that the residual is not exactly 0 guarantees a positive estimator for *σ*^2^. In the nonlinear case (10), this is not possible, and to guarantee existence of minimizers we have to obtain some conditions for model validity.

Let 𝒮_++_ denote the set of positive definite matrices, and for ***M*** ∈ 𝒮_++_ consider the function

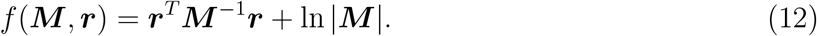

To connect the general functional form (12) with the problem (10), we specify functional domains for feasible ***r*** amd ***M***.

**Definition 1** (Domains). *We define domains* 𝒟_*r*_ *for* ***r*** *and* 𝒟_*M*_ *for* ***M*** *as follows:*

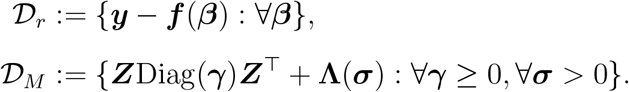

Using these definitions, we can write our objective as

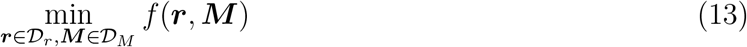

for *f* in (12). We also need to define the **level set** of a function.

#### Definition 2

(Level set). *The α-level set of a function f*: ℝ^*k*^ → ℝ, *denoted* ℒ_*f,α*_, *is defined by*

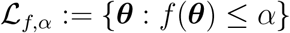

To guarantee the existence of estimators (***θ***) we make two assumptions.

#### Assumption 1.

*We assume that the image of* ***f*** *is closed. This implies that* 𝒟_*r*_ *is closed*.

#### Assumption 2.

*Denote the eigenvalue decomposition for any* ∀***M*** ∈ 𝒟_*M*_ *by* 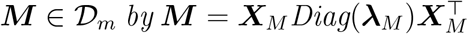. *For* ∀***r*** ∈ 𝒟_*r*_ *and* ∀***M*** ∈ 𝒟_*M*_, *we assume there exist α >* 0 *such that*

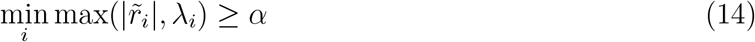

*where* 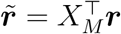 *and λ*_*i*_ *is the corresponding eigenvalue*.

Assumption 2 tells us that, after being pre-whitened by the covariance matrix, either the residual is bounded away from 0 or else the corresponding eigenvalue is bounded away from zero. These assumptions are necessary and sufficient for the existence of minimizers. Before we proceed, we make a simple remark about the meta-analysis case.

#### Remark 1.

*Assumption 2 is always satisfied for the case of meta-analysis, as long as all reported covariance matrices are positive definite*.

The remark follows because in the meta-analysis case, **Λ** is block diagonal, with blocks precisely the covariance matrices reported by studies. For many functions **f**, Assumption 1 is satisfied, including all linear and piecewise linear-quadratic functions. Functions that may violate these assumptions include logarithms and fractional functions, and complex models may therefore require further analysis. We can now prove the following lemma.

#### Theorem 1.

*Under Assumption 1 and Assumption 2, all level sets* ℒ_*f,α*_ *for f in* (13) *are bounded. Moreover, all level sets* ℒ_*f,α*_ *for f in* (13) *are closed*.

**Proof**

We first prove that

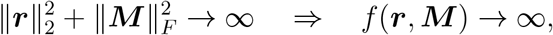

which implies bounded level sets. From (13) and the eigenvalue decomposition, we have

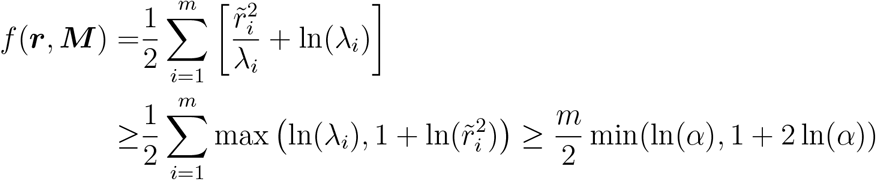

When 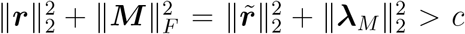 we know that there at least exists one 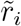 or *λ*_*i*_ such that 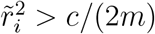 or 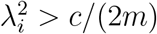, and

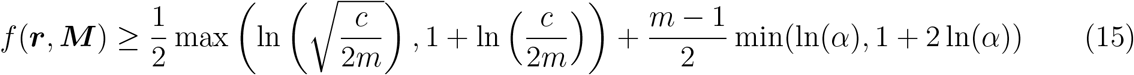

And we know when *c* → ∞, *f* (***r, M***) → ∞.

Next we prove that ℒ_*f,α*_ are also closed. Since *f* is the continuous function and 𝒟_*r*_ is closed from Assumption 1, we only need to show that the intersection between any ℒ_*c*_ and open boundary of 𝒟_*M*_, denoted as ℬ_*M*_, is empty. And we know that ℬ_*M*_ is contained in the space of positive indefinite matrices. For any 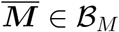 and any sequence ***M***^*i*^ ⊂ 𝒟_*M*_ approaching 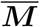, without loss of generality, we assume 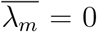, and we know that, 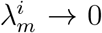 and the corresponding 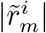 has to be stay above *α* (from Assumption 2). And from the inequality (15), we have

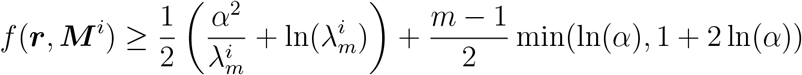

And we know 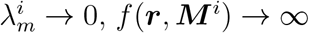, therefore there exists *N* such that *f* (***r, M*** ^*i*^) *> c* for all *i* ≥ *N*, and 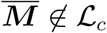.

We therefore have an immediate corollary.

#### Corollary 1

(Existence of minimizers). *Under Assumptions 2 and 1, the set of minimizers f is nonempty*.

By Theorem 1, the level sets of *f* are closed and bounded, hence compact. A continuous function assumes its minimum on a compact set, therefore the set of minimizers is nonempty. We can also use Theorem 1 to prove specific results about existence of minimizers for (10) under specific parametrizations, including meta-analysis and more standard longitudinal assumptions. Linear constraints will not fundamentally change the situation, so long as they allow any feasible solution, as is stated in the next corollary.

#### Corollary 2

(Effects of constraints). *As long as the intersection of domains* 𝒟_*r*_ *and* 𝒟_*M*_ *with the polyhedral set* ***Cθ*** ≤ ***c*** *is nonempty, minimizers exist under Assumptions 2 and 1. Moreover, constraints can be used to ensure Assumption 1, for specific cases, by ensuring that the intersection of the feasible region with* 𝒟_*r*_ *is closed*.

This follows immediately from the fact that the intersection of a compact set with any closed set remains compact.

Theorem 1 and following corollaries establish conditions for the existence of ***θ***(*w*), given assumptions 2 and 1. These assumptions must hold for all ***w*** in the capped simplex specified by the modeler through the *h* parameter, which roughly means that for any selection of *h* datapoints, the problem has to be well defined in the sense of assumptions 2 and 1.

The existence of global minimizers ***θ***(***w***) underpins the approach, which at a high level optimizers the value function ***v***(***w***) over the capped simplex to detect outliers. The optimization problem (10) has a nonconvex objective and nonlinear constraints. Nevertheless, we can specify the conditions under which ***v***(***w***) is a differentiable function, and we can find an explicit formula for its derivative. The result is an application of the Implicit Function Theorem to the Karush Kuhn Tucker conditions that characterize optimal solutions of (10). We summarize the relevant portion of this classic result presented by [Still, Theorem 4.4], and refer the reader to Craven [1984], Bonnans and Shapiro [2013] where these results are extended under weaker conditions.

#### Theorem 2

(Smoothness of the value function). *For a given* ***w***, *consider* ***θ***(***w***) *and let* I_0_ *denote the set of active constraints:*

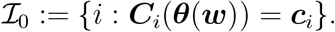

*Consider the extended Lagrangian function restricted to this index set:*

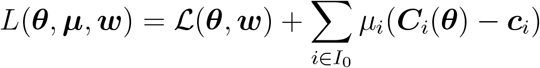

*Suppose that the following conditions hold:*

- *Stationarity of the Lagrangian: there exist multiplies* ***µ***(***w***) *with*

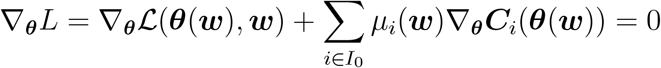
- *Linear independence constraint qualification*

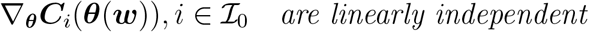
- *Either of the following conditions hold:*

-*Strict complementarity: number of active constraints is equal to the number of elements in* ***θ***, *and all µ*_*i*_(***w***) *>* 0.
-*Second order condition:* 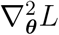 *is positive definite when restricted to the tangent space* 𝒯 *induced by the active constraints:*

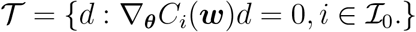

*Then the value function v*(***w***) *is differentiable, with derivative given by*

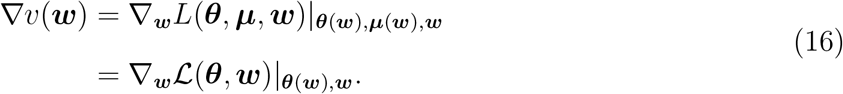

The second order condition above guarantees that ***θ***(***w***) is isolated (locally unique) minimizer [Still, Theorem 2.5].

Partially minimizing over ***θ*** reduces problem (10) to

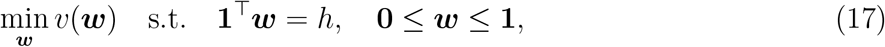

where *v*(***w***) is a continuously differentiable nonconvex function, and the constrained set is the (convex) capped simplex Δ_*h*_ introduced in the trimming section.

The high-level optimization over ***w*** is implemented using projected gradient descent:

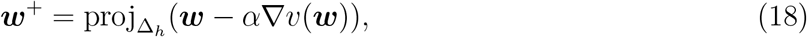

where 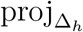 is the projection operator onto a set defined in the introduction. Since Δ_*h*_ is a convex set, the projection is unique.

Each update to ***w*** requires computing the gradient ∇*v*, which in turn requires solving for ***θ***; see (11). The explicit implementation equivalent to (18) is summarized in Algorithm 1. Projected gradient descent is guaranteed to converge to a stationary point in a wide range of settings, for any semi-algebraic differentiable function *v* (Attouch et al. [2013]).

**Algorithm 1.**
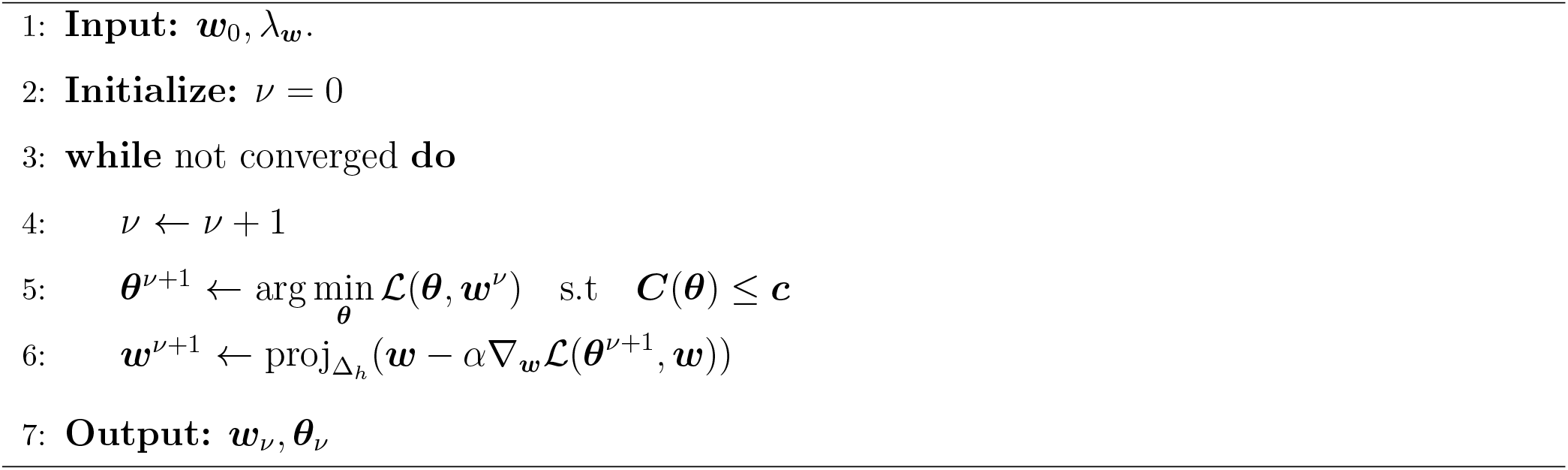
Projected gradient descent on the Value Function *v* of (11)

Step 5 of Algorithm 1 requires solving the constrained likelihood problem (10) with ***w*** held fixed. We solve this problem using `IPopt` [Wächter and Biegler, 2006], a robust interior point solver that allows both simple box and functional constraints. While one could solve the entire problem using `IPopt`, optimizing with simultaneously in ***θ*** and ***w*** turns the problem into a high dimensional nonsmooth nonconvex problem. Instead, we treat them differently to exploit problem structure. Typically ***θ*** is small compared to ***w***, which is the size of the data. On the other hand the constrained likelihood problem in ***θ*** is difficult while constrained value function optimization over ***w*** can be solved with projected gradient. We therefore iteratively solve the difficult small-dimensional problem using `IPopt`, while handling the optimization over the larger ***w*** variable through value function optimization over the capped simplex, a convex set.

### 2.6 Nonlinear Relationships using Constrained Splines

In this section we discuss using spline models to capture nonlinear relationships. The relationships most interesting to us are dose-response relationships, which allow us to analyze adverse effects of risk factor exposure (e.g. smoking, BMI, dietary consumption) on health outcomes. For an in-depth look at splines and spline regression see De Boor et al. [1978] and Friedman et al. [1991].

Constraints can be used to capture expert knowledge on the shape of such risk curves, particularly in segments informed by sparse data. Basic spline functionality is available in many tools and packages. The two main innovations here are (1) use of constraints to capture the shape of the relationship, explained in Section 2.6.2, and (2) nonlinear functions of splines, such as ratios or differences of logs, developed in Section 2.6.3. First we introduce basic concepts and notation for spline models.

#### 2.6.1 B-splines and bases

A spline basis is a set of piecewise polynomial functions with designated degree and domain. If we denote polynomial order by *p*, and the number of knots by *k*, we need *p* + *k* basis elements 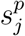, which can be generated recursively as illustrated in Figure 1.

**Figure 1:**
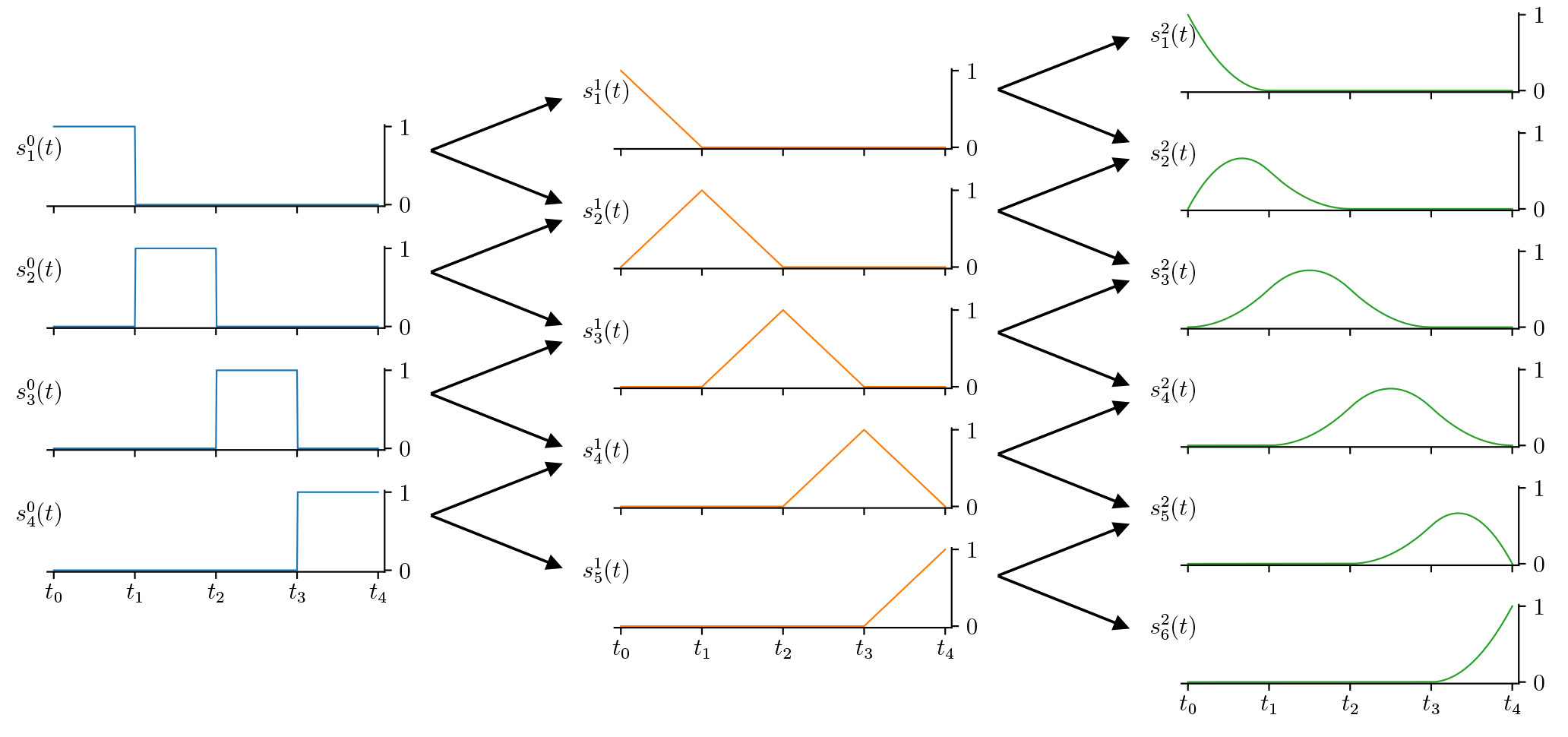
Recursive generation of bspline basis elements (orders 0, 1, 2). The points *t*_*i*_ in the figure are locations for knots, specified (in this case) to be equidistant. Once the spline bases are formed, they can then be evaluated at any points *t* in the domain, forming ***X*** as in (20).

Given such a basis, we can represent any nonlinear curve as the linear combination of the spline basis elements, with coefficients ***β*** ∈ ℝ^*p*+*k*^:

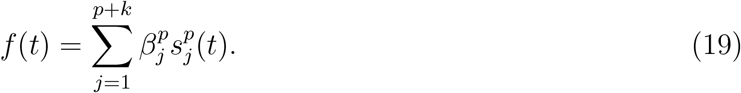

These coefficients are inferred by `LimeTr` analysis. A more standard explicit representation of (19) is obtained by building a design matrix ***X***. Given a set of *n*_*i*_ range values 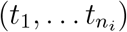 from study *i*, the *j*th column of 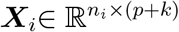 is given by the expression

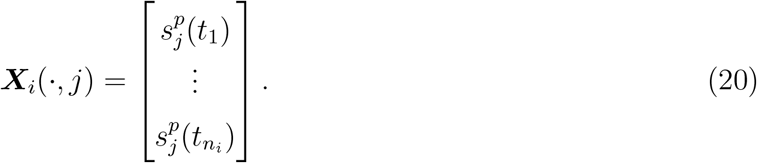

The model for observed data coming from (19) can now be written compactly as

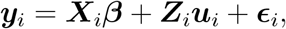

which is a special case of the main problem class (1).

#### 2.6.2 Shape constraints

We can impose shape constraints such as monotonicity, concavity, and convexity on splines. This approach was proposed by Pya and Wood [2015], who used re-formulations using exponential representations of parameters to capture non-negativity. The development in this section uses explicit constraints, which is simple to encode and extends to more general cases, including functional inequality constraints ***C***(***θ***) ≤ ***c***.

##### Monotonicity

Spline monotonicity across the domain of interest follows from monotonicity of the spline coefficients [De Boor et al., 1978]. Given spline coefficients ***β***, the curve *f* (*t*) in (19) is monotonically nondecreasing when

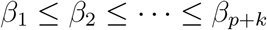

and monotonically non-increasing if

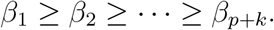

The relationship *β*_1_ ≤ *β*_2_ can be written as *β*_1_ − *β*_2_ ≤ 0. Stacking these inequality constraints for each pair (*β*_*i*_, *β*_*i*+1_) we can write all constraints simultaneously as

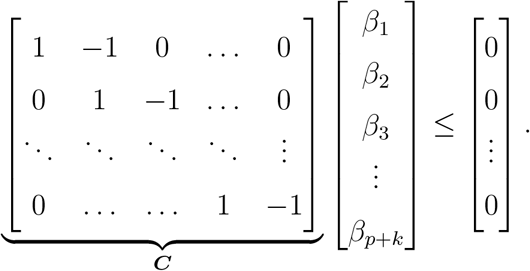

These linear constraints are a special case of the general estimator (10).

##### Convexity and Concavity

For any twice continuously differentiable function *f*: ℝ → ℝ, convexity and concavity are captured by the signs of the second derivative. Specifically, *f* is convex if *f* ^*″*^(*t*) ≥ 0 is everywhere, and concave if *f* ^*″*^(*t*) ≤ 0 everywhere. We can compute *f*^*″*^(*t*) for each interval, and impose linear inequality constraints on these expressions. We can therefore easily pick any of the shape combinations given in [Pya and Wood, 2015, Table 1], as well as imposing any other constraints on ***β*** (including bounds) through the interface of `LimeTr`.

#### 2.6.3 Nonlinear measurements

Some of the studies in Section 3.2 use nonlinear observation mechanisms. In particular, given a dose-response curve of the form (19), studies often report odds ratio of an outcome between exposed and unexposed groups that are defined across two intervals on the underlying curve:

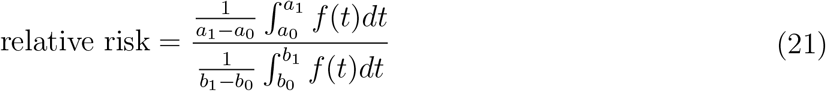

When *f* (*t*) is represented using a spline, each integral is a linear function of ***β***. If we take the observations to be the log of the relative risk for observation *j* in study *i*, this is given by

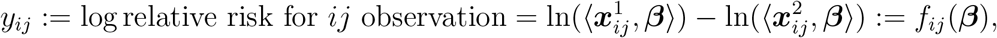

a particularly useful example of the general nonlinear term ***f***_*ij*_(***β***) in problem class (1). A set of examples of epidemiological models that arise from relative risk are discussed in Section 3.2.2.

### 2.7 Uncertainty Estimation

The `LimeTr` package modifies the parametric bootstrap [Efron and Tibshirani, 1994] to estimate the uncertainty of the fitting procedure. This strategy is necessary when constraints are present, and standard Fisher-based strategies for posterior variance selection do not apply [Cox, 2005].

The modified parametric bootstrap is similar to the standard bootstrap, but can be used more effectively for sparse data, e.g. when different studies sample sparsely across a dose-response curve. The approach can be used with any estimator (10).

In the linear Gaussian case, the standard bootstrap is equivalent to bootstrapping empirical residuals, since every datapoint can be reconstructed this way. When the original data is sparse, we modify the parametric procedure to sample *modeled* residuals. Having obtained the estimate 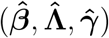, we sample model-based errors to get new bootstrap realizations 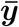.

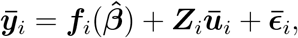

where 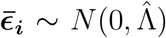 and 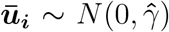. These realizations have the same structure as the input data. For each realization 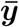, we then re-run the fit, and obtain *N* estimates 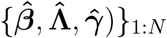. This set of estimates is used to approximate the variance of the fitting procedure along with any confidence bounds. The procedure can be applied in sparse and complex cases but depends on the initial fit 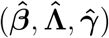. Its exact theoretical properties are outside the scope of the current paper and are a topic of ongoing research. We use N = 1000 in all of the numerical experiments in the next section. This significantly increases computational load, compared to a single fit. How to make the procedure more efficient (or develop alternatives) is another ongoing research topic.

## 3 Verifications

In this section we validate `LimeTr` on synthetic and empirical datasets. In Section 3.1 we show how `LimeTr` compares to existing robust packages on simple problems that can be solved by other available tools; see Table 1. We focus on robustness of the estimates to outliers, which is a key technical contribution of the paper.

In Section 3.2 we use the advanced features of `LimeTr` to analyze multiple datasets in public health, where robustness to outliers, and information communicated through constraints and nonlinear measurements all play an important role. These examples illustrate advanced `LimeTr` functionality that is not available in other tools. All examples in the paper are available online, along with a growing library of additional examples (see the Supplementary Materials section).

### 3.1 Validation Using Synthetic Data

Here we consider two common mixed effects models. First we look at meta-analysis, where the **Λ** term is known while (***β, γ***) are unknown. Then we look at a simple longitudinal case, where all three parameters are unknown, and **Λ** is modeled as *σ*^2^***I*** with unknown scalar *σ*^2^. In both of these cases, we compare the performance of `LimeTr` against several available packages. The simulated data is the same for both examples; only the model is different.

For the experiments, we generated 30 synthetic datasets with 10 observations in each of 10 studies (*n* = 100). The underlying true distribution is defined by *β*_0_ = 0, *β*_1_ = 5, *γ* = 6, and *σ* = 4, where *γ* is the standard deviation of the between-study heterogeneity and *σ* is the standard deviation of the measurement error. For the meta-analysis simulation, we assigned each observation a standard error of 4. The domain of the covariate *x*_1_ is [0, 10]. To create outliers, we randomly chose 15 data points in the sub-domain [6, 10] and offset them according to: 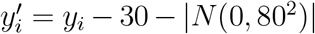. The setting of the simulation uses large outliers to show the robustness of the model to trimming. The larger the outliers, the larger the difference in performance between `LimeTr` and other tools. The behavior of trimming in the meta-analysis of real data is more clear in Section 3.2.

#### 3.1.1 Meta-analysis

We compared `LimeTr` to three widely-used packages that have some robust functionality: `metafor`, `robumeta`, and `metaplus`. The functionality developed in these packages differs from that of `LimeTr`. The `metafor` and `robumeta` packages refer to robustness in the context of the sandwich variance estimator, which makes the uncertainty around predictions robust to correlation of observations within groups. `metaplus` uses heavy-tailed distributions to model random effects, which potentially allows one to account for outlying studies but not measurements within studies.

Nonetheless, it is useful to see how a new package compares with competing alternatives on simple examples. We compared the packages to `LimeTr` in terms of error incurred when estimating ground truth parameters (*β*_0_, *β*_1_, *γ*), computation time, the true positive fraction (TPF) of outliers detected and the false positive fraction (FPF) of inliers incorrectly identified. If a threshold of 0.8 inliers is given to `LimeTr`, then outliers are exactly data points with an estimated weight *w*_*i*_ of zero, and those correspond to the largest absolute model residuals. To compare with other packages in terms of TPF and FPF, we identified the 20 data points with the highest residuals according to each packages’ fit. Table 2 and Figure 2 show the results of the meta-analysis simulation. The metrics are averages of 30 estimates from models fit on the synthetic datasets. `LimeTr` had lower absolute error in (*β*_0_, *β*_1_, *γ*), higher TPF, lower FPF and faster computation time than the alternatives.

**Table 2:** Results of Meta-Analysis Comparison. True values are *β*_0_ = 0, *β*_1_ = 5, *γ* = 6. Results show average estimates across realizations. The `LimeTr` package has much smaller absolute error in *β*_0_, *β*_1_ and in *γ* than other packages.

| Package | $\beta_0$ | $\beta_1$ | $\sqrt{\gamma}$ | TPF | FPF | Seconds |
| --- | --- | --- | --- | --- | --- | --- |
| Truth | 0 | 5 | 6 | — | — | — |
| **LimeTr** | **0.61** | **4.95** | **4.86** | **1.00** | **0.06** | **0.35** |
| robumeta | 10.92 | 0.03 | 37.2 | 0.68 | 0.12 | 6.54 |
| metaplus | 10.92 | 0.03 | 35.1 | 0.68 | 0.12 | 42.4 |
| metafor | 10.92 | 0.03 | 35.4 | 0.68 | 0.12 | 1.51 |

**Figure 2:**
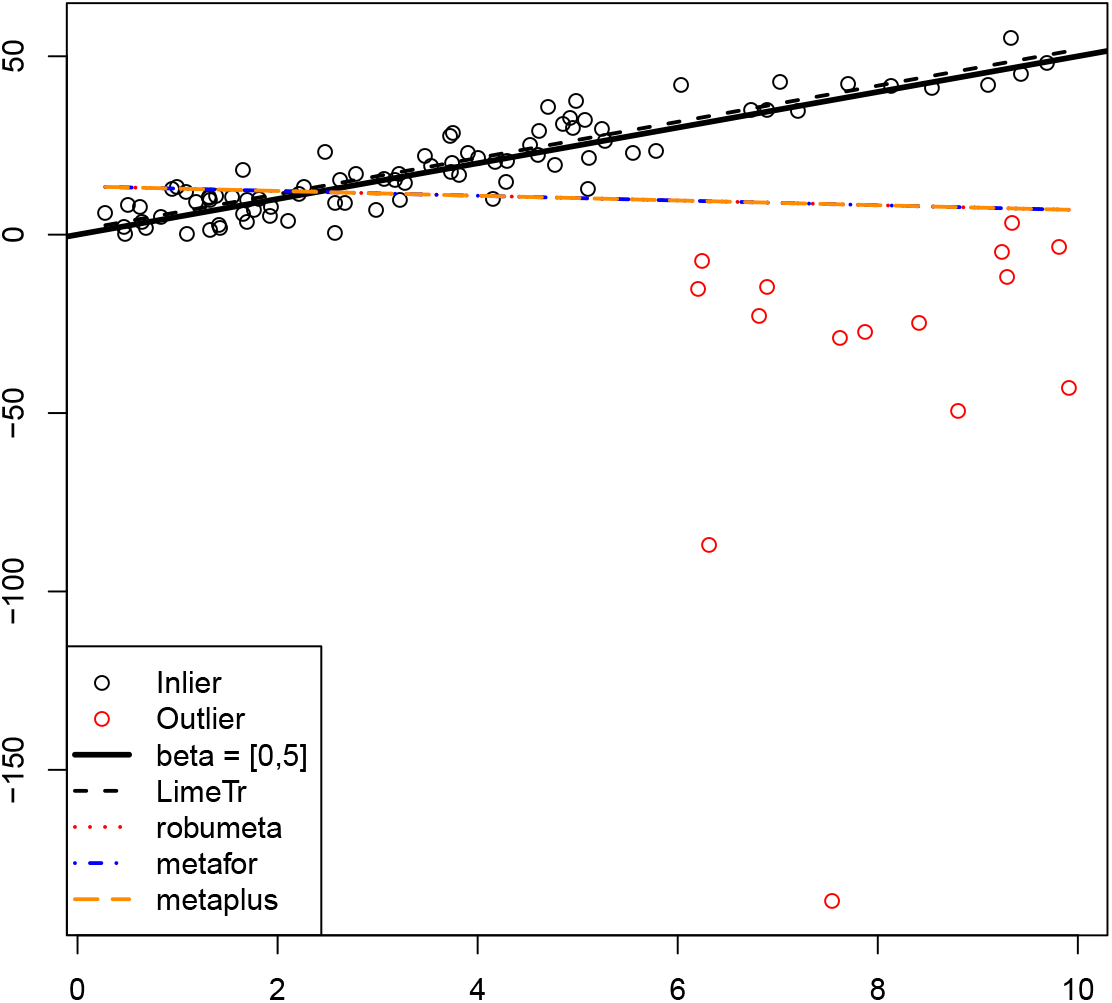
A representative instance of the experiment summarized in Table 2. True mechanism is shown using solid line; the true model is successfully inferred by the `LimeTr` package despite the outliers (red points).

**Figure 3:**
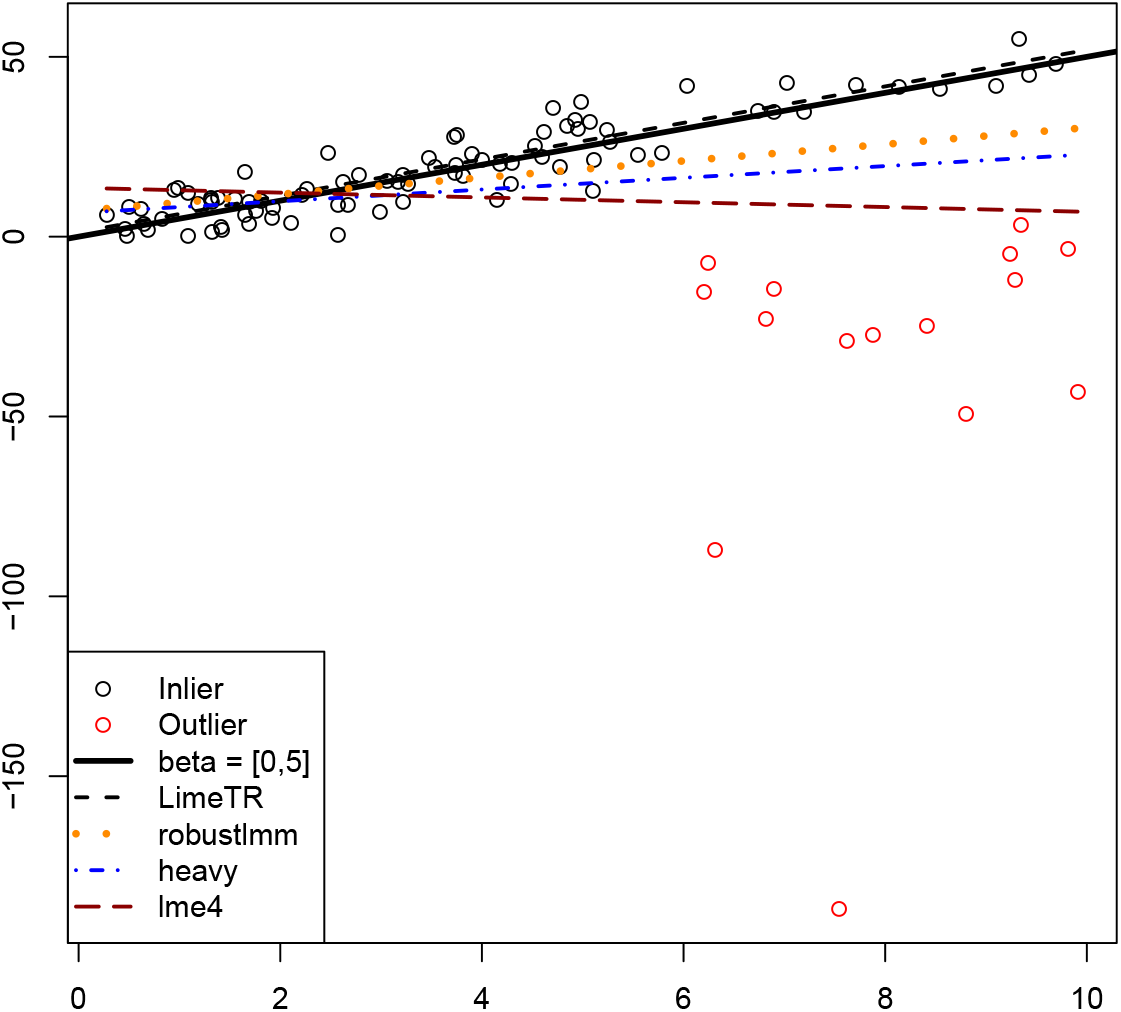
A representative instance of the experiment summarized in Table 3. `robustlmm` and `heavy` packages both estimate *β* better than `lme4`, likely because they use distributions with heavier tails. `LimeTr` outperforms all three alternatives.

#### 3.1.2 Longitudinal Example

Here we compare `LimeTr` to R packages for fitting robust mixed effects models. Rather than assuming that errors are distributed as Gaussian, the packages use Huberized likelihoods (`robustlmm`) and Student’s t distributions (`heavy`) to model contamination by outliers. `LimeTr` identifies outliers through the weights *w*_*i*_ in the likelihood estimate (3) that now captures simple longitudinal analysis. Specifically, **Λ** is no longer specified as in the meta-analysis case, but is instead parameterized through a single unknown error **Λ** = *σ*^2^***I*** common to all observations. We use the same simulation structure as in Section 3.1.1, now replacing observation-specific standard errors with random errors generated according to *N* (0, *σ*^2^). Since *σ*^2^ is now also an unknown parameter, we check how well it is estimated by all packages and report this error in Table 3.

**Table 3:** Results of Longitudinal Comparison. True values are *β*_0_ = 0, *β*_1_ = 5, *γ* = 6, *σ* = 4. Both `robustlmm` and the `heavy` package do better than the standard `lme4`. `LimeTr` more accurately estimates *β*_0_, *β*_1_, and *σ*. The `heavy` package estimates *γ* more accurately. However, estimated *γ* typically increases with a worse fit to the data, and heavy does not accurately estimate *β*.

| Package | $\beta_0$ | $\beta_1$ | $\sqrt{\gamma}$ | $\sigma$ | TPF | FPF | Seconds |
| --- | --- | --- | --- | --- | --- | --- | --- |
| Truth | 0 | 5 | 6 | 4 | — | — | — |
| LimeTr | **0.64** | **4.95** | 4.93 | **3.61** | **1.000** | **0.06** | 1.04 |
| robustlmm | 3.32 | 3.67 | **5.14** | 9.53 | 0.99 | 0.06 | 7.7 |
| heavy | 2.43 | 3.55 | 3.61 | NA | 0.95 | 0.07 | 0.16 |
| lme4 | 10.6 | 0.09 | 4.89 | 31.0 | 0.68 | 0.11 | **0.08** |

### 3.2 Real-World Case Studies

In this section we look at three real-data cases, all using meta-analysis. Across these examples, we show how trimming, dose-response relationships, and non-linear observation mechanisms come together to help understand complex and heterogeneous data.

#### 3.2.1 Simple Example: Vitamin A vs. Diarrheal Disease

The first example is a simple linear model that aims to quantify the effect of vitamin A supplementation on diarrheal disease. This is an important topic in global health, and we refer the interested reader to the Cochrane systematic review of the topic [Imdad et al., 2017]. In this example, we examine the influence outliers can have on inferences from a model, but we do not discuss in detail how to interpret the findings.

In this example, the dependent variable is the natural log of relative risk of incidence or mortality from diarrheal disease from 12 studies. Two of the studies report on both incidence and mortality, giving 14 total datapoints. No covariates are used. The model we consider is

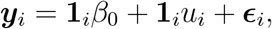

where ***y***_*i*_ are data sets reported from each study, **1**_*i*_ is a vector of 1’s of the same size as the number of observations for study *i*, _∈*i*_ ∼ *N* (**0, Σ**_*i*_) are associated standard errors, and *u*_*i*_ ∼ *N* (0, *γ*^2^) is a study-specific random effect, with *γ*^2^ accounting for between-study heterogeneity.

Figure 4 shows the results without trimming and with 10% trimming. We included data from 12 randomized control trials in the models, with a few studies having multiple observations. Without trimming, the estimated effect size is *β*_0_ = −0.15; with trimming it is three time smaller, *β*_0_ = −0.05. In the trimmed model, we observe all preserved points inside the funnel, indicating that there is no expected between-study heterogeneity. This is confirmed by the estimates – without trimming, between-study heterogeneity *γ*^2^ is estimated to be 0.0435, and after trimming two studies it is reduced to 7.14 × 10^*−*9^, nearly 0. Moreover, the potential outliers are unusual: while all other studies deal with ages 1 year or younger, the trimmed studies are among older ages, up to 10 years in one case and 3-6 years in the other. One of the trimmed studies was also conducted in slums, a non-representative population.

**Figure 4:**
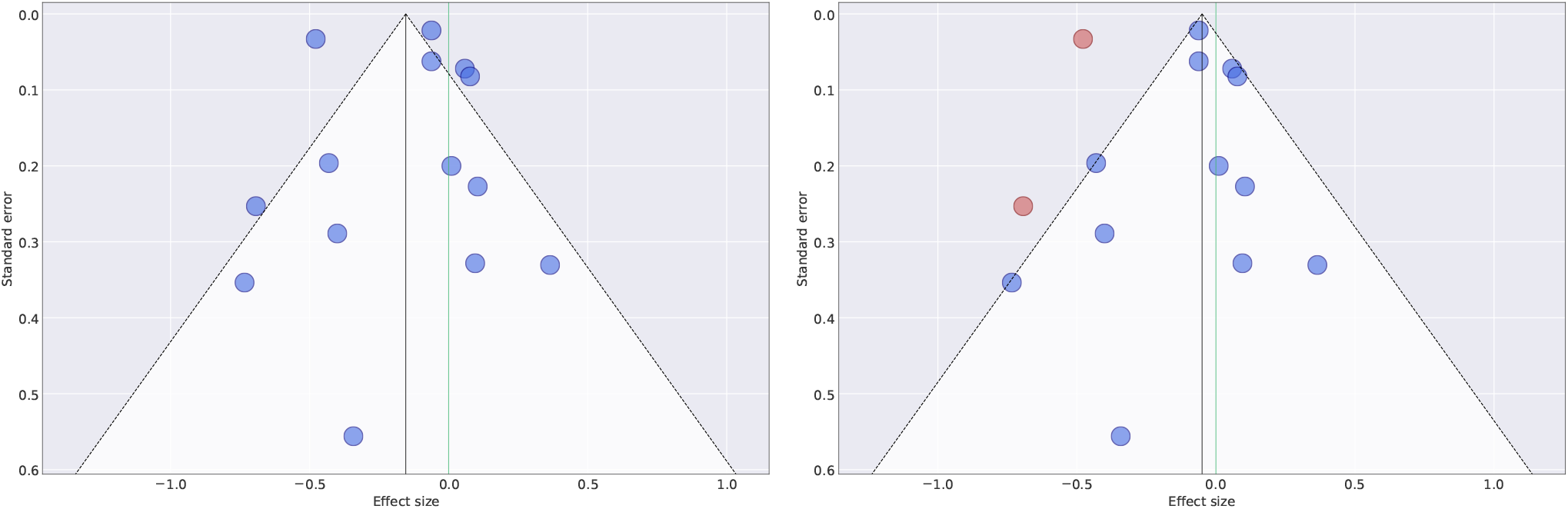
Results for Vitamin A vs. incidence and/or mortality rates for diarrheal disease, shown using the funnel plot (effect size vs. reported standard error). The left panel shows model without trimming, while the right panel shows the model with trimming. Trimming 10% of the studies shifts the effect closer to 0, making it less plausible that Vitamin A is protective, and identifies potential outliers (right panel, shown in red), which are able to hide more easily in the left panel.

#### 3.2.2 Spline Example: Smoking vs. Lung Cancer

##### log relative risk model

The log relative risk model is very common in the epidemiological context, and is often approximated by the log-linear relative risk model. A brief introduction to the nonlinear model was given in Section 2.6.3. Here we derive two models, and appropriate random effect specification, that can be used to analyze smoking and diet data.

In words, the log-relative risk model can be written as

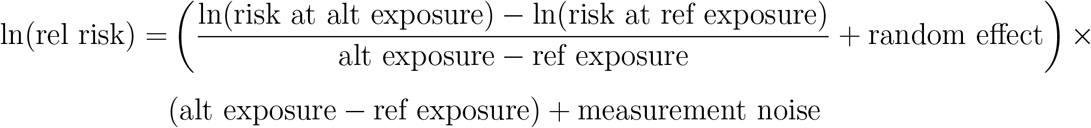

where the random effect is on the average slope and the measurement error in the log space. Instead of making the strong assumption that the log of the risk is a linear function of exposure, we use a spline to represent this relationship:

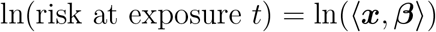

where ***x*** is the design vector at exposure *t* and the spline is parametrized by ***β***, see Section 2.6.1.

The correlation between smoking and lung cancer is indisputable [Gandini et al., 2008, Lee et al., 2012]. The exact nature of the relationship and its uncertainty requires accounting for the dose-response relationship between the amount smoked (typically measured in pack-years) and odds of lung cancer. We expect a nonlinear relationship between smoking and lung cancer, and the spline methodology described in Section 2.6.1 can be used.

The outcome here is the natural log of relative risk (compared to nonsmokers). The effect of interest is a function of a continuous exposure, measured in pack-years smoked. All studies compare different levels of exposure to non-smokers (exposure = 0), and we assume there is no risk for non-smokers (ln(risk at exposure 0) = 0). Since all datapoints share a common reference, we can simplify the relative risk model as follows:

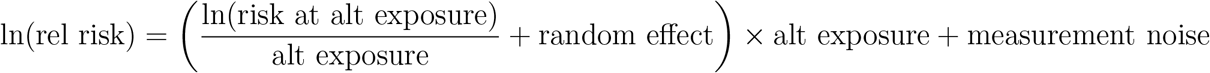

If we normalize risk of nonsmokers to 1, and use a spline to model the nonlinear risk curve, we have the explicit expression

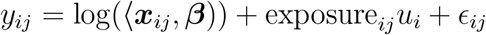

with *y*_*ij*_ the log relative risk, ***x***_*ij*_ computed using a spline basis matrix for exposure_*ij*_ (see Section 2.6.1), *u*_*i*_ the random effect for study *i*, and 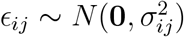 the variance reported by the *i*th study for its *j*th point.

To obtain the results in Figure 5 using `LimeTr`, interior knots were set at the 10th, 50th, and 80th percentiles of the exposure values observed in the data, corresponding to pack-year levels of 10, 30, and 55, respectively. We included 199 datapoints across 25 studies in the analysis, and again trimmed 10% of the data. Trimming in this case removes datapoints that are far away from the group (even considering between-study heterogeneity), as well as points that are closer to the mean but over-confident; these types of outliers are specific to meta-analysis.

**Figure 5:**
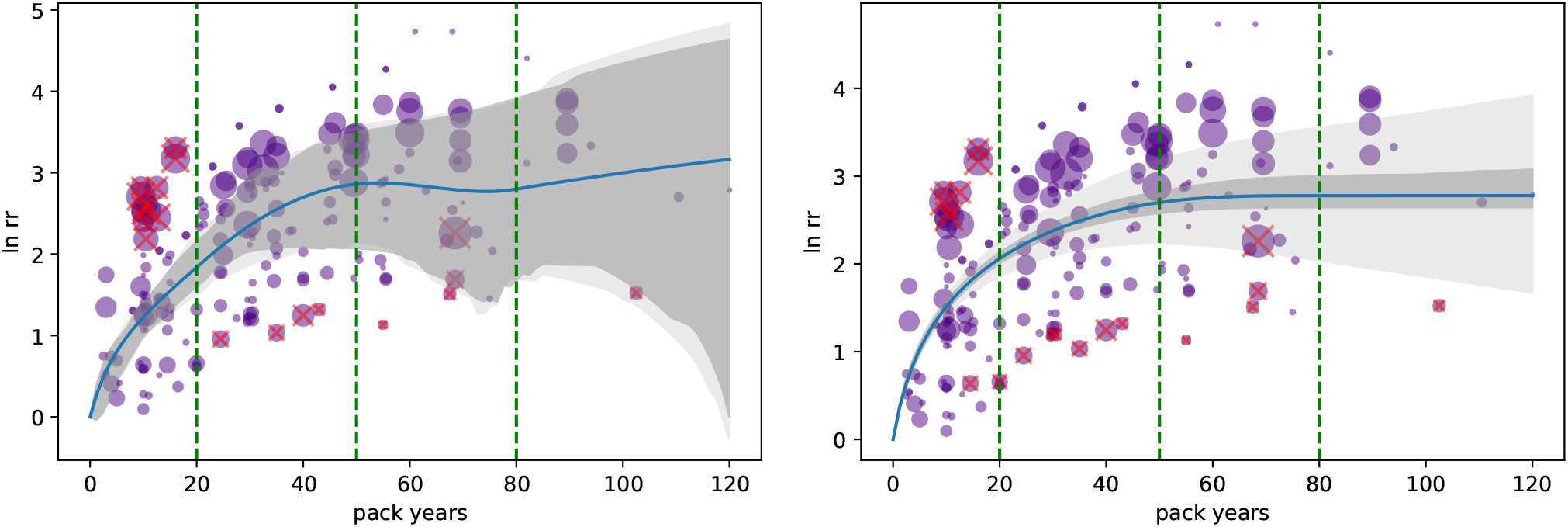
Modeling dose-response relationship between exposure (pack years) and log-odds of lung cancer. Left figure shows a cubic spline, while right figure shows a cubic spline with monotonically increasing risk and concavity constraints. The reference group is ‘nonsmokers’ in all studies, so taking nonsmoking risk at 1 we can plot the relative risk for each exposure group at its midpoint, and show the comparison to the estimated risk curve. The uncertainty of the mean is shown in dark grey, and additional uncertainty due to heterogeneity is shown in light gray. Constraints regularize the shape and decrease fixed effect uncertainty, but have higher estimated heterogeneity, whereas the more flexible model explains the data and has higher fixed effect uncertainty but lower heterogeneity. 10% trimming removes points that are far away from the mean dose-response relationship, as well as those moderately away from the mean but with very low reported standard deviation. Point radii on the graphs are inversely proportional to the reported standard deviations.

We also used multiple priors and constraints. First, we enforced that at an exposure of 0, the log relative risk must be equal to 0 (baseline: non-smokers) by adding direct constraint on *β*_0_. To control changes in slope in data sparse segments, we included a Gaussian prior of N(0, 0.01) on the highest derivative in each segment. Additionally, the segment between the penultimate and exterior knots on the right side is particularly data sparse, and is more prone to implausible behavior due to its location at the terminus, and we force that spline segment to be linear.

We show the unconstrained cubic spline in the left panel of Figure 5, and the constrained analysis that uses monotonicity and concavity of the curve in the right panel the same figure. The mean relationship looks more regular when using constraints, but the model cannot explain the data as well and so the estimate of heterogeneity is higher. On the other hand, the more flexible model has higher fixed effects uncertainty, and a lower estimate for between-study heterogeneity. The point sets selected for trimming are slightly different as well between the two experiments.

#### 3.2.3 Indirect nonlinear observations: red meat vs. breast cancer

The effect of red meat on various health outcomes is a topic of ongoing debate. In this section we briefly consider the relationship between breast cancer and red meat consumption, which has been systematically studied [Anderson et al., 2018]. In this section, we use data from available studies on breast cancer and red meat to show two more features of the `LimeTr` package: nonlinear observation mechanisms and monotonicity constraints.

The smoking example in Section 3.2.2 uses a direct observation model, since all measurement are comparisons to the baseline non-smoker group. This is not the case for other risk-outcome pairs. When considering the effect of red meat consumption, studies typically report multiplecomparisons between groups that consume various amounts of meat. In particular, all datapoints across studies are given to us a tuple: odds ratio for group [a, b] vs. group [c, d]. These datapoints are thus not measuring the spline directly, but are average slopes between points in log-derivative space. The observation model given by (21). As in the smoking example, we model the random effects on the average slopes of the log-relative risks, use a spline to represent the risk curve, and constrain the risk at zero exposure to be 1. In contrast to the smoking case, here we must account for both reference and alternative group definitions, so the observation model is given by

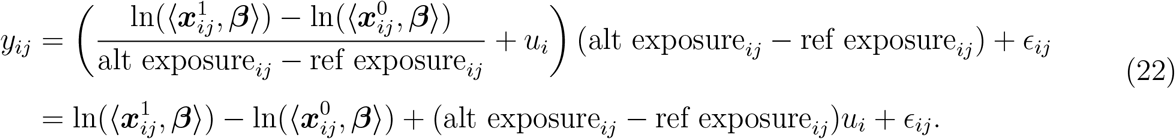

with *y*_*ij*_ the log relative risk,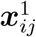 and 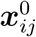 computed using spline basis matrix for alternative exposure_*ij*_ and reference exposure_*i*_, (see Section 2.6.1), *u*_*i*_ the random effect for study *i*, and 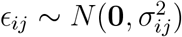 the variance reported by the *i*th study for its *j*th point.

`LimeTr` is the only package that can infer the nonlinear dose-response relationship in this example using heterogeneous observations of ratios of average risks across variable exposures. The meta-analysis to obtain the risk curve in Figure 6 integrates results from 14 prospective cohort studies^2^. An ensemble of spline curves was used rather than a single choice of knot placement. The cone of uncertainty coming from the spline coefficients is shown in dark gray in the left panel, with additional uncertainty from random effects heterogeneity shown in light gray.

**Figure 6:**
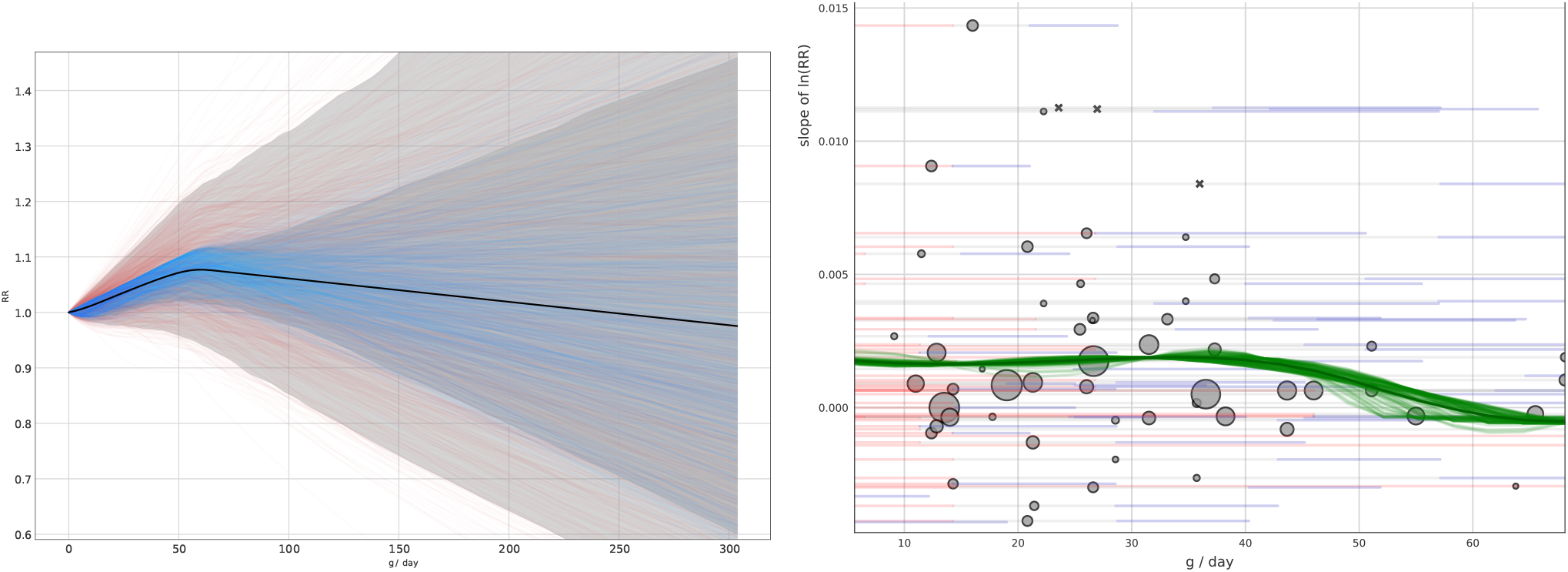
Spline fit obtained from observation model (22). The left panel shows the inferred dose response mechanism from indirect observations, across multiple spline knot placement options. Uncertainty from the spline coefficient fits is plotted in dark gray, with additional uncertainty coming from the random effects shown in light gray. Observations are shown in the right panel. The midpoint between the reference and alternative intervals is displayed as a point, with the reference and alternative exposures plotted in blue and red.

The right panel of Figure 6 shows the data fit. Each point represents a ratio between risks integrated across two intervals, so in effect on four points that define these intervals. We choose to plot the point at the midpoint defined by these intervals for the visualization. As in other plots, the radius of each point is inversely proportional to the standard deviation reported for it by the study. The derivatives of each spline curve fit are plotted in green so that they can be compared to these average ratios on log-derivative space.

After studying the relationship and the data in Figure 6, it is hard to justify forcing a monotonic increase in risk. However, to test the functionality of `LimeTr`, we add this constraint and show the result in Figure 7. Our conclusions about the relationship and its strength would change in this example if we impose such shape constraints, unlike the example that compares smoking and lung cancer.

**Figure 7:**
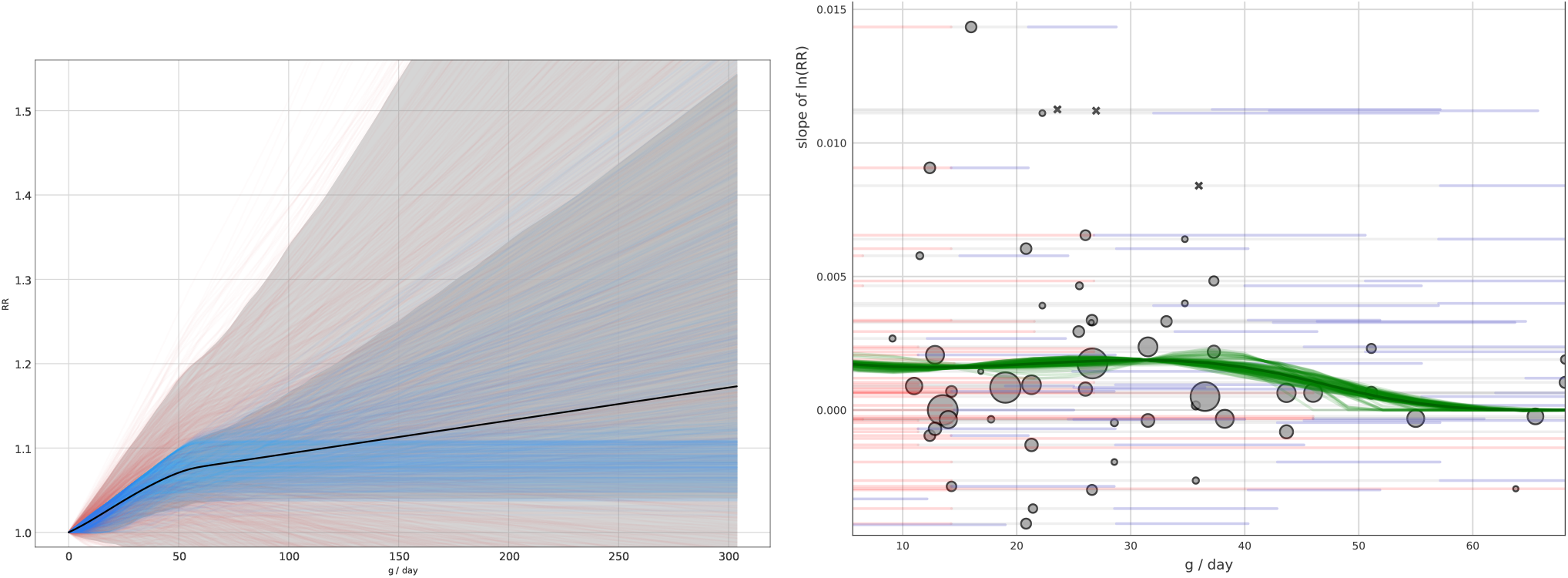
Adding monotonicity constraints to example in Figure 6.

## 4 Conclusion

We have developed a new methodology for robust mixed effects models, and implemented it using the `LimeTr` package. The package extends the trimming concept widely used in other robust statistical models to mixed effects. It solves the resulting problem using a new method that combines a standalone optimizer `IPopt` with a customized solver for the value function over the trimming parameters introduced in the reformulation. Synthetic examples show that `LimeTr` is significantly more robust to outliers than available packages for meta-analysis, and also improves on the performance of packages for robust mixed effects regression/longitudinal analysis.

In addition to its robust functionality, `LimeTr` includes additional features that are not available in other packages, including arbitrary nonlinear functions of fixed effects ***f***_*i*_(***β***), as well as linear and nonlinear constraints. In the section that uses empirical data, we have shown how these features can be used to do standard meta-analysis as well as to infer nonlinear dose-response relationships from direct and indirect observations of complex nonlinear relationships.

## SUPPLEMENTAL MATERIALS

### LimeTr package

Python package `LimeTr` containing code to perform robust estimation of mixed effects models, for both meta-analysis and simple longitudinal analysis. Available online through github: https://github.com/zhengp0/limetr

### Experiments

Set of script files available online to produce simulated data and run `LimeTr` and third party code: https://github.com/zhengp0/limetr/tree/paper/experiments

- `Settings.R`: R code to Specifies folder structure and simulation parameters.
- `functions.R`: R code for auxiliary functions for simulating data and aggregating results.
- `0_create_sim_data.R`: R code to create simulated data.
- `1_limetr.py`: Python code to run `LimeTr`.
- `2_metafor.R`: R code to run `metafor` package
- `3_robumeta.R`: R code to run `robumeta` package
- `4_metaplus.R`: R code to run `metaplus` package
- `5_lme4.R`: R code to run `lme4` package
- `6_robustlmm.R`: R code to run `robustlmm` package
- `7_heavy.R`: R code to run `heavy` package

## Footnotes

1 Breakdown refers to the percentage of outlying points which can be added to a dataset before the resulting M-estimator can change in an unbounded way. Here, outliers can affect both the outcomes and training data.

2 Variation in risk here is much lower than for smoking, so we show relative risk instead of log relative risk.

